# A critical contribution of a sparse neuronal ensemble in the amygdala central nucleus to extinction

**DOI:** 10.1101/2022.03.02.481339

**Authors:** Belinda P. P. Lay, Eisuke Koya, Bruce T. Hope, Guillem R. Esber, Mihaela D. Iordanova

## Abstract

Adaptive behaviour critically depends on the delicate and dynamic balance between acquisition and extinction memories. Disruption of this balance, particularly when the extinction memory loses control over behaviour, is the root of treatment failure of maladaptive behaviours such as substance abuse or anxiety disorders. Understanding this balance requires a better understanding of the underlying neurobiology and its contribution to behavioural regulation. Here, we used Daun02 in *Fos-lacZ* transgenic rats to delete extinction-recruited neuronal ensembles in BLA and CN and examine their contribution to behaviour in an appetitive Pavlovian task. Deletion of extinction-activated ensembles in CN but not BLA impaired the retrieval of extinction and increased activity in the BLA. The disruptive effect of deleting these CN ensembles was enduring as it hindered further extinction learning, and promoted greater levels of behavioural restoration across opposing levels of the response scale seen in spontaneous recovery and reinstatement. Our data indicate that the initial extinction-recruited CN ensemble is critical to the acquisition-extinction balance, and that greater behavioural restoration does not mean weaker extinction contribution. These findings provide a novel avenue for thinking about the neural mechanisms of extinction and in developing treatments for cue-triggered appetitive behaviours.

---

A curious property of learning systems is that while it may take a single trial to acquire a new association (e.g. tone→ food), no amount of extinction trials (tone→ nothing) will suffice to unlearn it. Indeed, after extensive extinction training, the mere passage of time (Pavlov, 1927; Rescorla, 1997; Rescorla and Heth, 1975) or an encounter with the outcome by itself (Rescorla and Heth, 1975) can cause behaviour to slip back under the control of the original acquisition memory. This acquisition bias is widely taken to imply that extinction training promotes not the erasure of the original memory, but the formation of a new inhibitory memory that survives alongside that of acquisition and competes for behavioural control. While the endurance of acquisition memories can be advantageous if environmental conditions revert to their initial state, the frailty of extinction memories presents a problem for behavioural modification (e.g. Conklin and Tiffany, 2002; Marlatt, 1990), where undesirable actions often prove resistant to extinction or prone to relapse. For this reason, it is essential to elucidate the role of extinction in this competition.

At the neural level, one reason why extinction is transient may be because it recruits its own neuronal ensemble that competes with that of acquisition (Herry et al., 2008; Lacagnina et al., 2019) as opposed to modifying the original acquisition memory. Neuronal ensembles within the basolateral (BLA) and central (CN) nuclei of the amygdala are suggestive of such an interaction as distinct neuronal subpopulations respond to cues that predict either the presence or absence of appetitive or aversive outcomes (Calu et al., 2010; Esber and Holland, 2014; Esber et al., 2012; Haubensak et al., 2010; Herry *et al*., 2008; Holland and Gallagher, 1993; Iordanova et al., 2016; Kim et al., 2017; Li et al., 2013; Roesch et al., 2010; Tye and Janak, 2007). Targeting CN and BLA extinction-recruited ensembles will answer some fundamental questions regarding their contribution to this competition. First and foremost, we can determine whether a small subset of anatomically localized extinction-activated neurons is *necessary* for extinction-governed behaviour. If so, we can further establish whether, once recruited, this specific ensemble has a long-lasting involvement in learning during future extinction episodes, making extinction neurobiologically constrained. Lastly, we can determine whether extinction ensembles lose control over behaviour when the balance between acquisition and extinction is upset in favour of the former.

To provide insight into these questions, we employed the Daun02/*Fos-lacZ* inactivation method, which permits targeted deletion of function-specific neuronal ensembles. This method allows strongly activated neurons to express β-galactosidase (β-gal) under the control of the *Fos* promoter (Koya et al., 2009). When the prodrug Daun02 is microinjected, β-gal catalyses the conversion of Daun02 into daunorubicin, which reduces neuronal excitability (Barker et al., 2017) and induces apoptosis in behaviourally-activated neuronal ensembles (Pfarr et al., 2015). Critically, this technique leaves neighbouring cells intact (Cruz et al., 2013). Using this targeted approach, we show that deletion of extinction-activated neuronal ensembles in CN but not BLA disrupts extinction retrieval, hinders further extinction learning, and promotes the original acquisition memory as suggested by signature restoration phenomena (i.e., spontaneous recovery and reinstatement). To our knowledge, we provide the first evidence that a small and localized subset of neurons can have a substantive and lasting effect on appetitive extinction-governed behaviour. Our findings showcase the anatomical vulnerability of extinction memories and shed new light on the pervasiveness of relapse in reward disorders.

## RESULTS

### CN but not BLA neurons are preferentially recruited during extinction

We characterised BLA and CN neurons activated during extinction. Rats learned to discriminate between a target cue paired with pellets and a non-reinforced control cue (Figures 1A and 1B; see Supplemental for data). After discrimination training, half the rats received extinction of the target cue (Extinction group), while the other half continued to receive nonreinforced presentations of the control cue (Control group; Figure 1B). This training was followed by fluorescent immunohistochemistry to label Fos, β-gal, protein kinase C delta (PKCδ), and somatostatin (SOM; Figures 1C-1E) in the BLA (Lee et al., 2016) and CN (Adke et al., 2021; Kim *et al*., 2017). Within the BLA (Figure 1D; Supplemental Figure 1A), there were no differences between extinction and controls in the number of Fos-immunoreactive neurons, nor in the number of double-labelled neurons (Fos + β-gal, Fos + PKCδ, Fos + SOM, max *t*(10) = 1.81, *p* = 0.10, 95% CI [-0.24, 2.33]). Within the CN (Figure 1E; Supplemental Figure 1B), extinction led to more Fos immunoreactive neurons (*t*(10) = 3.10, *p* = 0.011, 95% CI [0.50, 3.08], d = 1.79), and more Fos and β-gal co-localisation (*t*(10) = 3.46, *p* = 0.006, 95% CI [0.71, 3.28], d = 2.00) compared to controls. There were no differences in Fos + SOM overlap between extinction and control (*t*(10) = 0.15, *p* = 0.88, 95% CI [-1.20, 1.37]), but fewer Fos + PKCδ-expressing neurons were activated in extinction compared to controls (*t*(10) = 3.04, *p* = 0.012, 95% CI [-3.04, -0.47], d = 1.76). The degree of PKCδ and SOM expression as well as co-labelling of Fos and β-gal and Fos and PKCδ in the CN is consistent with those reported by others (Bossert et al., 2011; Fanous et al., 2012; Funk et al., 2016; Venniro et al., 2018). These findings show that while extinction recruits similar number of CN and BLA neurons, CN neuron recruitment is extinction-specific. Moreover, SOM+ neurons were not preferentially activated during extinction in the CN or BLA, whereas PKCδ+ neurons in the CN were activated by the non-reinforced control cue but not during early extinction/omission detection (see Discussion).

**Figure 1.**
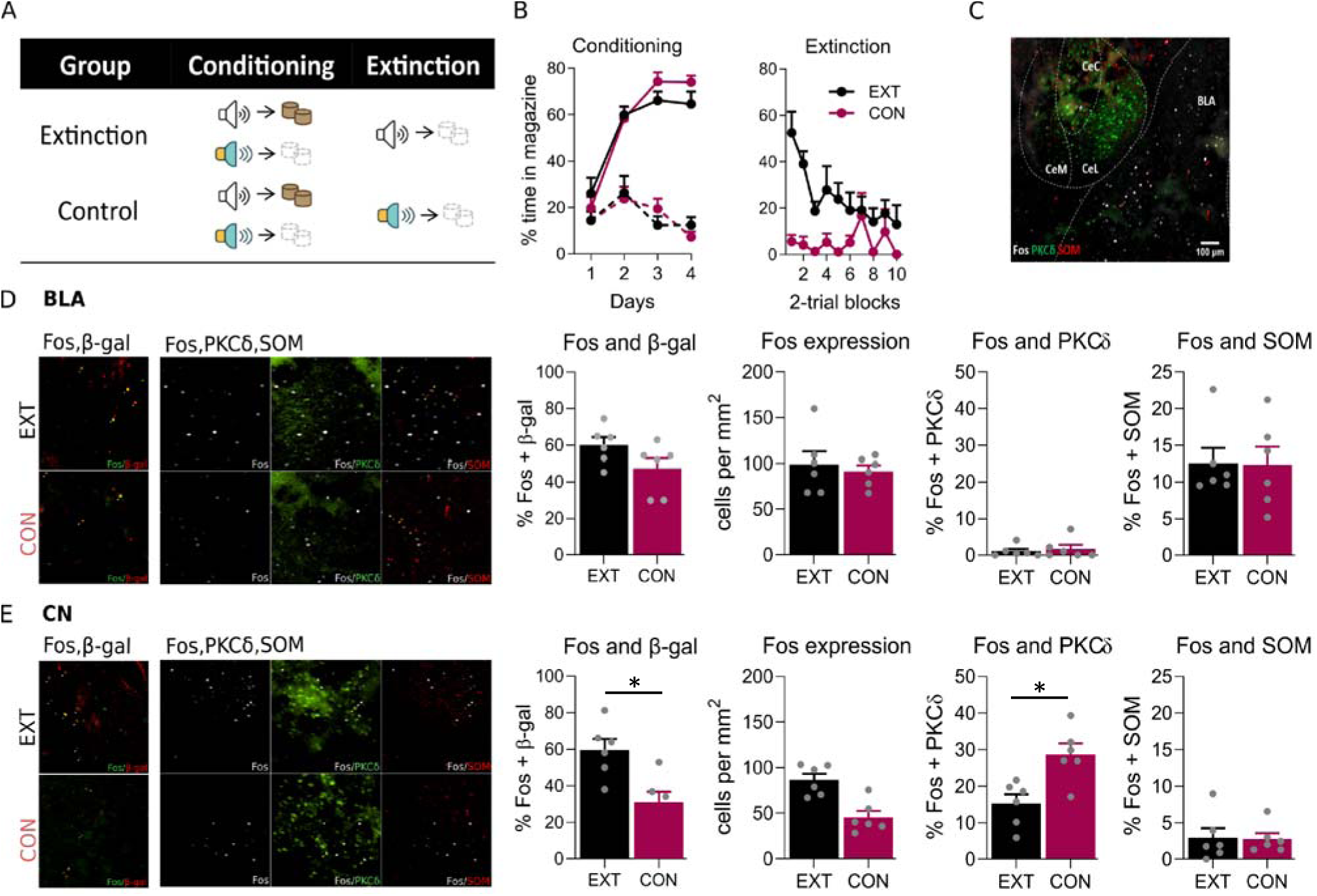
CN but not BLA neurons are preferentially recruited during extinction. A) Behavioural protocol. B) Behavioural data represent mean (+SEM) percent time spent in the magazine during the target cue for Conditioning and Extinction. Extinction (EXT, black), n = 6, and control (CON, burgundy), n = 6. C) Representative photomicrographs of Fos, PKCδ and SOM expression in the CN and BLA. Scale bar, 100 µm. Representative photomicrographs and mean percent colocalization (+SEM) of Fos+β-gal, Fos, Fos+PKCδ, or Fos+SOM double-labelled neurons during extinction (top) and control (bottom) in the D) BLA and E) CN. Arrows denote co-localisation of Fos with β-gal, PKCδ or SOM.

### Extinction-activated neurons in the BLA are not critical for extinction expression

While there was no differential activation of BLA neurons during extinction compared to the control, the overall level of activation was similar to that seen in the CN. Therefore, to determine whether these extinction-activated BLA neurons are necessary for extinction (see Figure 2A for design), we targeted these function-specific neurons using Daun02 inactivation (see Figure 2B for placements). Rats were trained to discriminate between two auditory cues (e.g. target→pellet, control→nothing, Figure 2C; see Supplemental for data). After discrimination training, rats received non-reinforced presentations of the target or control cue during extinction (Figure 2C; see Supplemental for data), followed by Daun02 or vehicle infusions into the BLA. All rats were then tested for conditioned responding to the target cue to assess whether extinction retrieval depended on the deleted extinction-activated BLA neurons. Our findings show that this deletion had no effect on extinction retrieval. (Figure 2C right panel; main effect of training, *F*_(1, 31)_ = 18.84, *p* < 0.001, 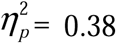, 95% CI [-1.17, -0.31], d = 1.45; no main effect of drug and no interaction, *Fs*<1).

**Figure 2.**
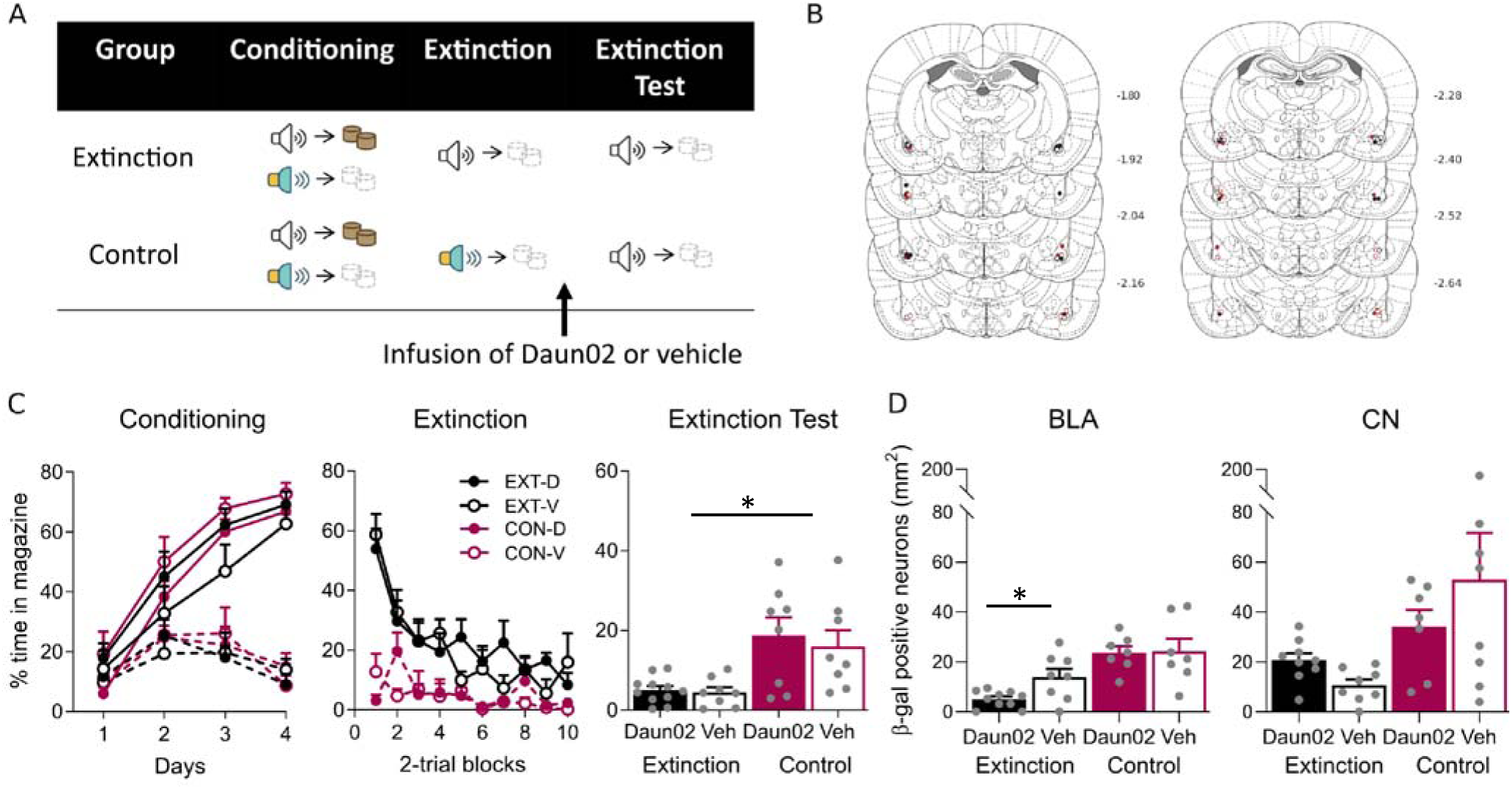
Extinction-activated neurons in the BLA are not critical for extinction expression. A) Behavioural protocol. B) BLA cannula placements, distances in millimetres from bregma. C) Mean (+SEM) percent time spent in the magazine during the target cue for Conditioning, Extinction, and Extinction Test. Extinction-Daun02 (EXT-D, filled black), n = 11, extinction-vehicle (EXT-V, open black), n = 8, control-Daun02 (CON-D, filled burgundy), n = 8, and control-vehicle (CON-V, open burgundy), n = 8. Solid lines refer to responding to the reinforced, target cue and the dashed lines refer to responding to the non-reinforced, control cue. D) Mean (+ SEM) number of β-gal positive neurons in the BLA (left) and CN (right) to the extinction cue on test. Extinction-Daun02, n = 10 (BLA) and 9 (CN), extinction-vehicle, n = 8, control-Daun02, n = 7, and control-vehicle, n = 7 (BLA) and 8 (CN).

Quantification of β-gal positive neurons in the BLA confirmed the effectiveness of the Daun02 manipulation (Figure 2D left panel). This was evident in the fewer β-gal positive neurons in the Daun02 compared to the vehicle extinction groups (*F*_(1, 28)_ = 4.79, *p* = 0.037, 95% CI [-2.01, -0.066], d = 1.19), but similar counts in the control groups (*F*<1). In the same brains, the total number of β-gal positive neurons in the CN region were also quantified (Figure 2D right panel). There was no effect of drug on the expression of β-gal positive neurons in the CN in either the extinction nor control groups (max*F*_(1, 28)_ = 1.65, *p* = 0.21, 95% CI [-1.73, 0.40]).

### Extinction-activated neurons in the CN are critical for extinction expression

To determine whether extinction-activated CN neurons were critical for extinction retrieval, we targeted these function-specific neurons using Daun02 inactivation (for design and placements see Figures 3A and 3B). Rats received identical discrimination and extinction training to that described previously (Figures 3C; see Supplemental for data), followed by Daun02 or vehicle infusions into the CN. To determine whether extinction retrieval depended on extinction-activated CN neurons, all rats were tested for conditioned responding to the target cue. Silencing extinction-recruited CN neurons disrupted extinction retrieval (Figure 3C; extinction-Daun02 vs. extinction vehicle: *F*_(1, 57)_ = 9.39, *p* = 0.003, 95% CI [0.17, 0.79], d = 1.03) leaving responding in those animals similar to that in controls (*F*_(1, 57)_ = 1.12, *p* = 0.30, 95% CI [-0.59, 0.18]). Importantly, this disruption in extinction was specific to the extinction of the target cue as intra-CN infusion of Daun02 after *reinforced* presentations of the target cue did not disrupt responding at test (*t*(19) = 1.11, *p* = 0.28, 95% CI [-0.43, 1.40], Figure 3C right panel).

**Figure 3.**
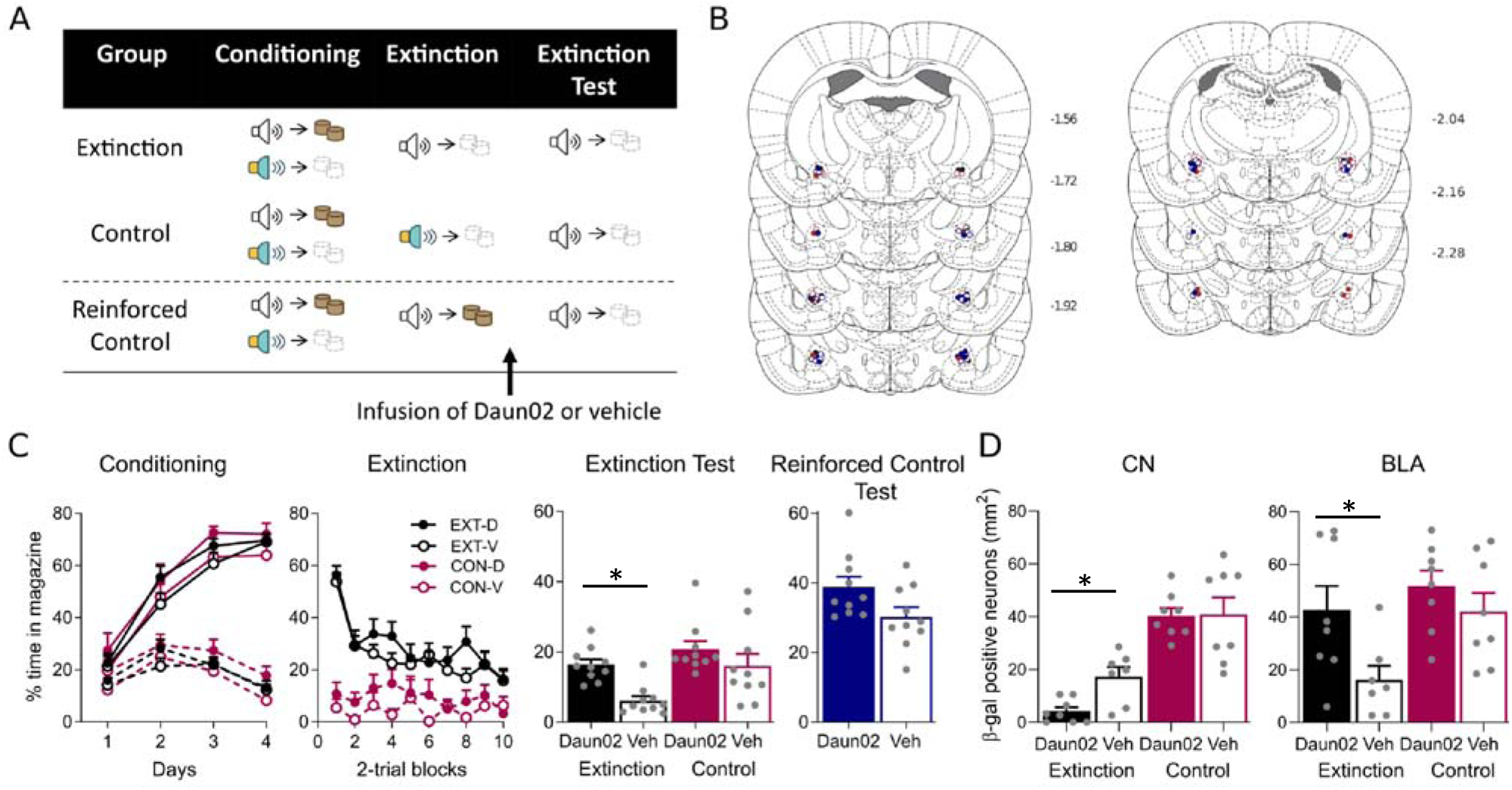
Extinction-activated neurons in the CN are critical for extinction expression. A) Behavioural protocol. B) CN cannula placements, distances in millimetres from bregma. C) Mean (+SEM) percent time spent in the magazine during the target cue for Conditioning, Extinction, Extinction Test, and Reinforced Control Test. Extinction-Daun02 (EXT-D, filled black), *n* = 20, extinction-vehicle (EXT-V, open black), *n* = 21, control-Daun02 (CON-D, filled burgundy), *n* = 10, control-vehicle (CON-V, open burgundy), *n* = 10. Reinforced control-Daun02 (filled blue), *n* = 11, and reinforced control-vehicle (open blue), *n* = 10. Solid lines refer to responding to the reinforced, target cue and the dashed lines refer to responding to the non-reinforced, control cue. D) Mean (+ SEM) number of β-gal positive neurons in the CN (left) and BLA (right) to the extinction cue on test. Extinction-Daun02, *n* = 8, extinction-vehicle, *n* = 7, control-Daun02, *n* = 8, and control-vehicle, *n* = 8.

Daun02 deletion in the CN was also effective as seen in the fewer β-gal positive neurons in the Daun02 extinction group compared to the vehicle extinction group (Figure 3D left panel; *F*_(1, 27)_ = 5.01, *p* = 0.034, 95% CI [-2.22, -0.10], d = 1.74) but, as expected, not in the controls (*F*<1). Interestingly, there were more β-gal positive neurons in the BLA in the extinction-Daun02 group compared to the extinction-vehicle group (*F*_(1, 27)_ = 6.77, *p* = 0.015, 95% CI [0.28, 2.41], d = 1.29; Figure 3F right panel) but not in the controls (*F*<1). This suggests the possibility that high levels of responding indicative of disrupted extinction may be driven by activation of BLA neurons. Indeed, in the groups that did not show extinction, number of BLA β-gal positive neurons positively correlated with the level of conditioned responding on the first trial-block of the test (extinction-Daun02: *r*(6) = 0.70, *p* = 0.05; combined control groups: *r*(14) = 0.57, *p* = 0.02).

### The original extinction-activated neurons in the CN regulate subsequent extinction learning and spontaneous recovery

To determine if the CN neurons that were recruited initially during extinction are also necessary for further extinction learning, a subset of extinction-Daun02 and extinction-vehicle rats were retained following the test from the study above and tested again 24hrs (re-test) and nine days later (spontaneous recovery, Figure 4A). During the 24hrs re-test, responding was higher in the extinction-Daun02 group compared to the extinction-vehicle group (Figure 4C; *F*_(1, 22)_ = 6.08, *p* = 0.022, 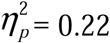, 95% CI [0.04, 0.43], d = 0.98), indicating that further extinction learning accruing during this test partly relied on the originally recruited CN ensemble (for placements see Figure 4B). However, by the end of this test session responding in all groups had extinguished to the same point (*F*_(1, 22)_ = 0.548, *p =* 0.47, 95% CI [-0.55, 1.15]). Critically, when responding was tested eight days later, spontaneous recovery was greater in the Daun02 compared to the vehicle rats (*F*_(1, 14)_ = 6.61, *p* = 0.022, 95% CI [0.05, 0.55], d = 1.22; Figure 4D). Thus, deletion of an extinction-recruited ensemble in CN led to a persistent disruption in extinction responding despite additional training as well as greater spontaneous recovery.

**Figure 4.**
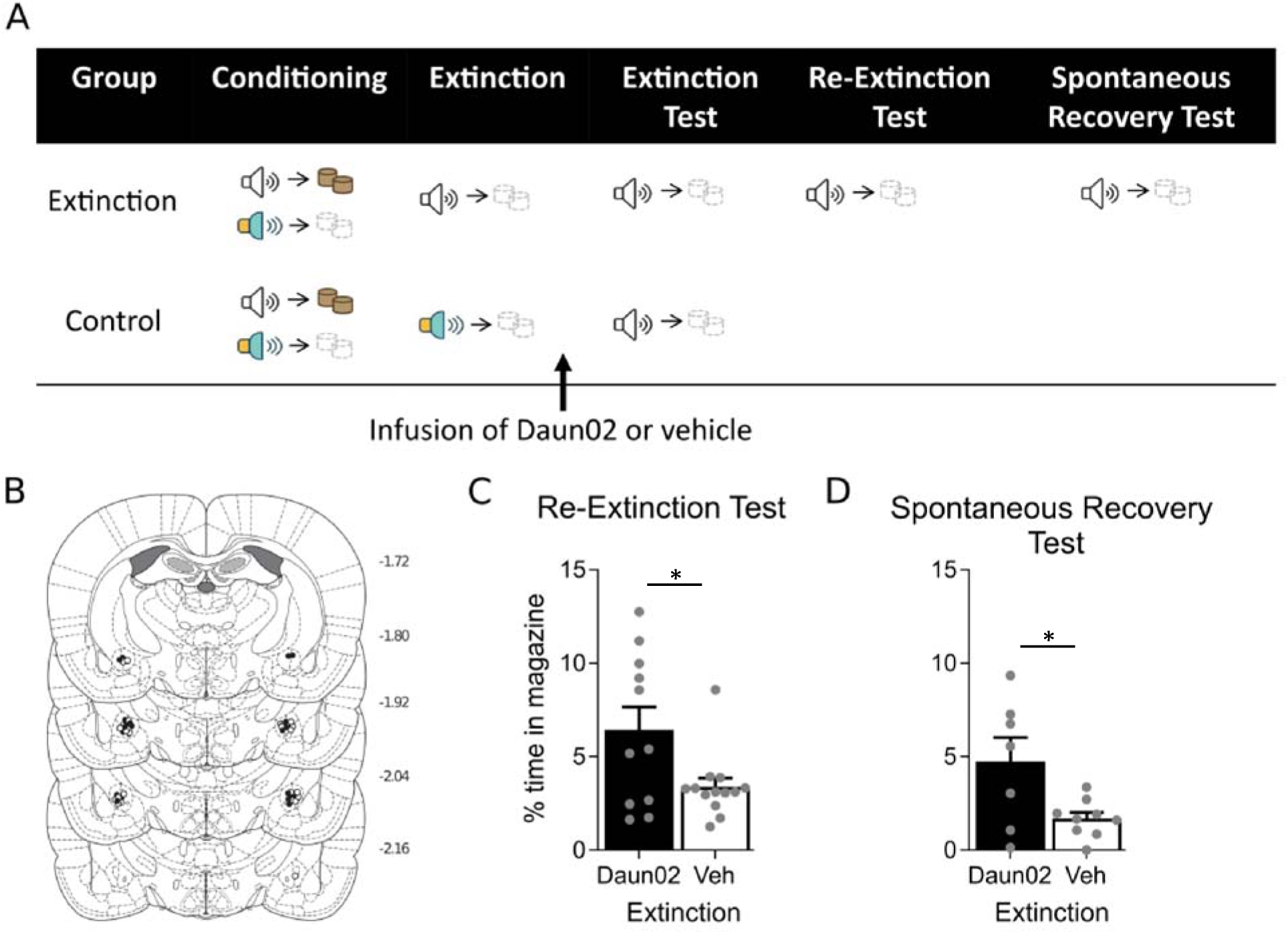
The original extinction-activated neurons in the CN regulate subsequent extinction learning and spontaneous recovery. A) Behavioural protocol for Re-extinction and Spontaneous Recovery Tests. B) CN cannula placements, distances in millimetres from bregma. Mean (+SEM) percent time spent in the magazine during the target cue for C) Re-Extinction Test and D) Spontaneous Recovery Test. Extinction-Daun02 (EXT-D, filled black), n = 11 (Re-Extinction) and n = 7 (Spontaneous Recovery), extinction-vehicle (EXT-V, open black), n = 13 (Re-Extinction) and n = 9 (Spontaneous Recovery).

### Deletion of extinction-activated neurons in the CN enhances reinstatement

Behavioural restoration methods shift the balance between acquisition and extinction in favour of the former, but they can be arranged to do so to different degrees. For example, the incomplete level of spontaneous recovery (as reported here) suggests that both acquisition and extinction memories continue to exert some behavioural control. Indeed, deleting the extinction-recruited CN neurons facilitated behavioural restoration. If, however, greater restoration indicates greater control of the acquisition memory over behaviour, then the deletion of the extinction-recruited CN neurons should have little if any effect. An alternative holds that the extinction memory does not cease to to influence responding when behavioural restoration is high, but this influence might be obscured by the greater contribution of the acquisition memory. To assess the contribution of extinction under conditions of strong behavioural restoration, we used a reinstatement design which consisted of unsignaled exposure to the outcome prior to test (see Figures 5A and 5B for design and placements, respectively). We gave rats discrimination training followed by extinction training (Figure 5C, see Supplemental for data) before injecting them with either Daun02 or vehicle in the CN. Prior to testing with the target cue, a subset of rats received exposure to the outcome (pellets; Reinstatement-Daun02 and Reinstatement-vehicle; Figure 5C right panel), which reinstated responding to the extinguished cue in the vehicle group compared to a non-reinstatement control (*F*_(1, 22)_ = 10.77, *p* = 0.003, 95% CI [0.17, 1.47], d = 2.63). Critically, silencing extinction-activated CN neurons after extinction training augmented the reinstatement effect (Reinstatement-Daun02 vs. Reinstatement-vehicle: *F*_(1, 22)_ = 9.66, *p* = 0.015, 95% CI 0.11, 1.21], d = 1.36). This finding indicates that extinction-recruited CN neurons continue to influence behaviour even under conditions that make that influence difficult to the detect.

**Figure 5.**
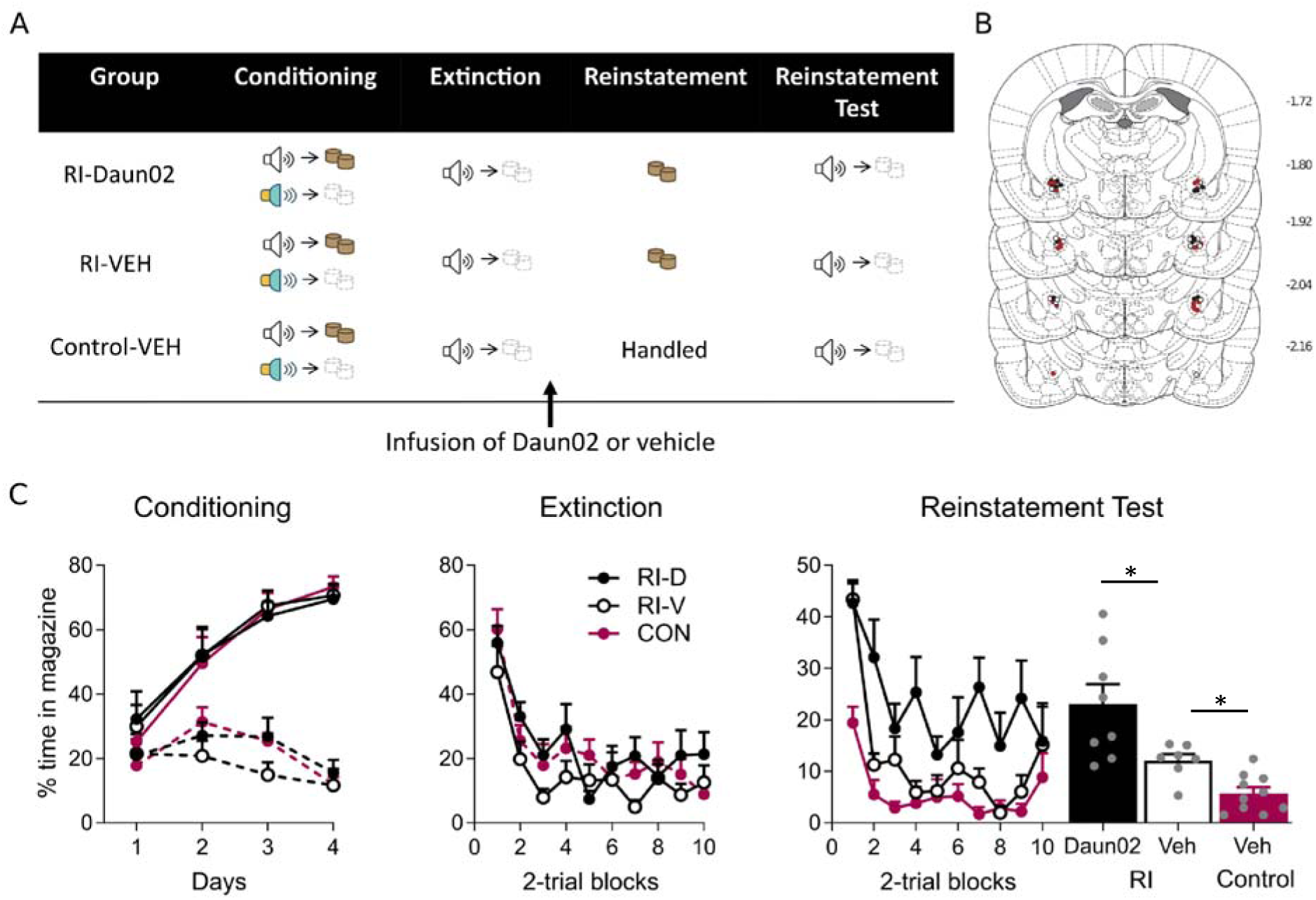
Deletion of extinction-activated neurons in the CN enhances reinstatement. A) Behavioural protocol for Reinstatement. B) CN cannula placements, distances in millimetres from bregma. C) Mean (+SEM) percent time spent in the magazine during the target cue for Conditioning, Extinction, and Reinstatement Test. Reinstatement-Daun02 (RI-D, filled black), n = 8, reinstatement-vehicle (RI-V, open black), n = 7, and control-vehicle (CON-V, filled burgundy), n = 10. Solid lines refer to responding to the reinforced, target cue and the dashed lines refer to responding to the non-reinforced, control cue.

## DISCUSSION

Here, we disrupted the balance between extinction and acquisition by deleting extinction-recruited neurons in the BLA and CN. Consequently, we found that CN but not BLA extinction ensembles are critical for extinction retrieval and further extinction learning. In addition, deletion of these ensembles enhanced the behavioural control exerted by the initial acquisition memory following the passage of time (spontaneous recovery) and outcome re-exposure (reinstatement). Our results shed new light on the relative contribution of amygdala nuclei to extinction in reward learning, and indicate that decision-making after extinction hinges on a delicate balance between acquisition- and extinction-specific neuronal ensembles.

By uncovering that extinction relies on a subset of neurons that are anatomically localized and activated following a single extinction episode, we show that extinction memories are sparsely represented in the brain. This sparsity likely signifies a weaker neurobiological footing relative to that of acquisition, making extinction easily disrupted when environmental conditions change. In other words, limited neuronal recruitment positions them to be easily outcompeted by other populations. Strikingly, this originally recruited extinction ensemble has an *enduring* involvement in regulating extinction despite additional training. This finding is particularly important because it shows that the sparsity of neuronal recruitment during extinction is not (fully) compensated by other ensembles as learning continues. Thus, the vulnerability of extinction to disruption is compounded by the long-lasting engagement of a specific subset of neurons. Therefore, our data provide fundamental neurobiological insight into a central feature of extinction; that is, its high incidence of relapse.

Despite the sparsity of this extinction ensemble, our data imply that it continues to compete with the memory of acquisition even under conditions that favour the latter. This is evident in the lower level of responding in the vehicle group (where the ensemble is intact), than in the Daun02 group following spontaneous recovery and even outcome exposure (reinstatement), two scenarios marked by an acquisition bias. However, the fact that extinction was observed at all in the Daun02 group suggests that additional extinction ensembles were present within the CN either because the neuronal deletion was incomplete or because additional neurons were recruited during subsequent extinction training, albeit insufficiently to compensate for the original ensemble. Of course, other ensembles elsewhere in the brain likely contributed to extinction, yet once again failed to offset the loss of the original CN ensemble.

One important question raised by our findings concerns the nature of the information encoded by the extinction-recruited ensembles in CN. On the one hand, the deleted CN neurons may contribute to storing a cue→no-outcome association formed during extinction in order to compete with the original cue→outcome association. Eliminating such neurons would thus grant the latter association the upper hand in behavioural control. On the other hand, the deleted ensemble might be critical for restoring attention to the cue in the face of unexpected reward omission and increasing the rate of extinction learning elsewhere (Holland and Gallagher, 1993; Schiffino and Holland, 2016). Although our data are not able to dissociate between these possibilities, both accounts predict that the behavioural scales should be persistently tilted in favour of acquisition, consistent with our findings.

Our data further reveal an important dissociation between the BLA and CN in extinction. Unlike with CN, deleting BLA extinction-activated ensembles did not disrupt extinction retrieval at test. Yet BLA could still play a role in the balance of acquisition and extinction memories. Deleting extinction-activated CN ensembles led to a concomitant increase in β-gal expression in BLA, which positively correlated with an enhanced responding to the previously extinguished cue (i.e., disrupted extinction). This correlation was also present in the control (non-extinction) groups, suggesting that the BLA might promote acquisition-based responding (but see Hatfield et al., 1996). Thus, the balance between acquisition and extinction may depend on the interaction between BLA and CN neuronal ensembles.

Studies in fear suggest that one potential mediator of BLA-CN interaction is the locus coeruleus (LC; Cedarbaum and Aghajanian, 1978; Giustino and Maren, 2018; Valentino and Van Bockstaele, 2008; Van Bockstaele et al., 1998). Neuropeptide corticotropin-releasing factor (CRF) expressing neurons in the CN promote the activity of noradrenergic neurons in the LC that project to BLA (McCall et al., 2015; see also Prouty et al., 2017; Schwarz et al., 2015; Van Bockstaele *et al*., 1998 see Iordanova, Yau, McDannald, & Corbit, 2021 for review of LC in extinction), rendering conditioned responding (in fear) more persistent in general (Giustino et al., 2020; Sanford et al., 2017; see also Uematsu et al., 2017) and in the face of extinction (Abiri et al., 2014; Giustino *et al*., 2020; Sanford *et al*., 2017; see also Uematsu *et al*., 2017). Therefore, it is possible that extinction might hinder the ability of CRF-neurons in CN to boost the activity of BLA acquisition ensembles (via LC activation, Giustino and Maren, 2018), limiting the latter’s influence over behaviour.

Another finding that merits consideration is the lower level of activity of PKCδ-expressing neurons to the extinction compared with the control cue after a single extinction episode. This result may seem paradoxical in view of a literature linking PKCδ neurons to behavioural inhibition across a variety of settings, including fear suppression following extinction (Ciocchi et al., 2010; Haubensak *et al*., 2010), suppression of feeding (Cai et al., 2014), and social reward-induced reduction in incubation of drug craving (Venniro *et al*., 2018). However, our observations may in fact be consistent with that literature. In our study, the control cue initially received extended training in the absence of the outcome and on the background of target cue→outcome pairings. This would result in the involvement of PKCδ neurons as behavioural inhibition develops to the control cue, such that when that cue is subsequently presented these neurons increase their level of activity. The low level of activity in these neurons to the extinction cue may reflect their lack of involvement in the early detection of outcome omission that drives extinction learning. Rather, PKCδ neuronal activity may be elevated once extinction and the resulting behavioural inhibition are well-established (e.g. on day 2).

To conclude, our findings indicate that extinction relies on the pervasive influence of a relatively small subset of anatomically localized neurons, in the absence of which extinction training is ineffectual. There are two ways to think about the implications of these findings for maladaptive behaviour. On the one hand, the recruitment of such a lean neural substrate would explain why extinction memories are so fragile in the face of environmental change, be it in the laboratory or the clinical setting. It further suggests that recalcitrant disorders characterized by the long-term inefficacy of extinction treatments might result from an even leaner recruitment of extinction ensembles. If so, then an extinction-centered approach to behavioural modification might be less fruitful than one based on reducing the influence of acquisition through disruption in reconsolidation (e.g. Duvarci et al., 2008; Nader et al., 2000). On the other hand, our findings identify a potential locus for future neural interventions based on enhancing the recruitment of extinction-activated ensembles. An optimistic note is found in the long-lasting effects a localized ensemble can have on behaviour, which suggest the possibility of leveraging such an ensemble long after the initial extinction treatment. Our data thus offer a novel way of thinking about the underlying causes of resistance to extinction-based therapies and the pervasiveness of relapse.

## METHODS

### Subjects

One hundred and ninety-nine male Sprague Dawley *Fos-LacZ* transgenic rats (RRID:RRRC_00865) weighing between 350-470g were obtained in-house from the *Fos-LacZ* breeding colony at Concordia University. Littermates were counterbalanced across groups in each experiment to avoid a litter effect. Rats were pair-housed in a standard clear cage (44.5 cm x 25.8 cm x 21.7 cm) containing a mixture of sani-chip and corncob bedding. The boxes were kept in an air-conditioned colony room maintained on a 12-hr light-dark cycle (lights off at 8:00 am). Food and water were available ad libitum prior to surgery and during recovery, and thereafter food was restricted to 10g/rat/day. All experimental procedures were in accordance with the approval granted by the Canadian Council on Animal Care and the Concordia University Animal Care Committee.

#### Sex

The present studies use only males despite starting the investigation with both sexes. This is because females take longer to learn from extinction (Delamater et al., 2017; see also Lay et al., 2020), making it impossible to use the Daun02 inactivation procedure following a single day of extinction training.

### Surgery and Drug Infusion

Before behavioural training and testing, rats were implanted with bilateral guide cannulae in the BLA or CN. They were anaesthetized with isoflurane gas and then mounted on a stereotaxic apparatus (David Kopf Instruments). They were then treated with a subcutaneous injection of rimadyl (5mg/kg; Pfizer, Kirkland, QC) immediately upon placement in the stereotaxic frame. Twenty-two-gauge single-guide cannulae (Plastics One) were implanted through holes drilled in both hemispheres of the skull above the BLA (AP: -2.4 mm, ML: ±4.7 mm, DV: -7.6 mm) or CN (AP: -2.4 mm, ML: ±4.0 mm, DV: -7.2 mm). DV coordinates were taken relative to skull surface at site of implantation. The guide cannulae were secured to the skull with four jeweller’s screws and dental cement. A dummy cannula was kept in each guide at all times except during microinjections. Rats were allowed six days to recover from surgery, during which time they were handled, weighed, and given an oral administration of 0.5 ml solution of cephalexin daily.

Daun02 (2µg/0.5µl) or vehicle was infused bilaterally into the BLA or CN by inserting a 28-gauge single injector cannula into the guide cannula. The injector cannula was connected to a 10-µL Hamilton syringe attached to an infusion pump (Harvard Apparatus). The injector cannula projected an additional 1 mm ventral to the tip of the guide cannula. A total volume of 0.5 µl was delivered to both sides at a rate of 0.25 µl/min, and drug delivery was monitored with the progression of an air bubble in the infusion tube. The injector cannula remained in place for an additional 2 min after the infusion to allow for drug diffusion before its complete removal. Immediately after the infusion, the injector was replaced with the original dummy cannula. One day before infusions, all rats were familiarised with this procedure by removing the dummy cannula and inserting the injector cannula to minimise stress the following day.

### Drugs

A DNA synthesis inhibitor, Daun02 (A3352, ApexBio), was dissolved in 5% dimethyl sulfoxide (DMSO; Bioshop), 6% Tween 80 (Bioshop), and 89% 0.1M phosphate buffered saline (PBS, pH 7.4) to obtain a final concentration of 4ug/ul. The vehicle solution consisted of 5% DMSO, 6% Tween 80, and 89% 0.1M phosphate buffered saline (PBS, pH 7.4).

### Behavioural Apparatus

#### Stimuli

Two 10-s auditory cues were used in the experiments. One was a 10-Hz, 75-dB mechanical clicker located on the outside left panel of the behavioural chamber; the other was a 72-dB white-noise delivered through a loud speaker located outside the behavioural chamber. The cues were controlled via a Med-Associates program. The background noise in the chamber was 48-50 dB. Sound intensity was measured using a digital sound level meter (Tenma, 72-942) located inside the chamber and placed flat in the middle of the grid floor facing the magazine. The cues were fully counterbalanced. Chocolate-flavoured sucrose pellets (Product# F07256, Bio-Serv, Flemington, NJ) served as reinforcers.

#### Experimental chambers

Behavioural procedures were conducted in 16 operant-training chambers, each measuring 31.8 cm in height x 26.7 cm in length x 25.4 cm in width (Med Associates, St. Albans, VT, USA). The modular left and right walls were made of aluminium, and the back wall, front door, and ceiling were made of clear Perspex. Their floors consisted of stainless-steel rods, 4 mm in diameter, spaced 15 mm apart, center to center, with a tray below the floor. Illumination of each chamber was provided by a red house-light mounted 1 cm below the ceiling on the centre panel of the right wall. Each chamber was enclosed in a ventilated sound attenuating cabinet. The back wall of each cabinet was equipped with a camera connected to a monitor located in another room of the laboratory where the behaviour of each rat was observed by an experimenter. Stimuli were presented through Med Associates software on a computer located outside the experimental room. Eight of the chambers had checkered or spotted wallpaper on the door and each wall with the exception of the back wall to allow for video viewing. Instead, the back wall of the holding cabinet was covered in either checkered or spotted wallpaper. The chambers consisting of the checkered or spotted wallpaper are referred to as the pattern chambers and the remaining eight chambers are referred to as the bare chambers. The pattern chambers were cleaned with 4% almond-scented solution (President’s Choice Black Label) and the bare chambers with 10% acetic acid solution. The chamber type was counterbalanced across groups.

### Behavioural Procedures

#### Experiment 1: CN but not BLA neurons are preferentially recruited during extinction

##### Magazine Training

One day prior to the start of behavioural training, all rats received 20 chocolte-flavoured sucrose pellets in their home cage to reduce neophobia during training. Twenty-four hours later, rats were placed in the conditioning context and received magazine training (40-min session). During this session, sucrose pellets were delivered into the magazine at one-minute intervals (total of 40 pellets) in order to train the rats to approach the magazine. No stimuli were presented during this phase. At the end of this and all remaining training sessions, the chambers (walls, door, grid floor, and tray) were cleaned (0.5% acetic acid solution in the bare chambers; 1% almond solution in the pattern chambers) after removal of each rat.

##### Phase 1 Conditioning

On days 1 to 4, rats were trained to discriminate between two auditory cues (white-noise or clicker) such that one (the target cue) was reinforced, whereas the other (control) was not. Rats were placed into the conditioning context, and after a 2 min adaptation period, they received ten presentations of the target stimulus followed by the delivery of two sucrose pellets at the 8^th^ and 9^th^ second of cue delivery, and ten non-reinforced presentations of the control stimulus. Stimuli were 10-s in duration and were counterbalanced across all rats. Stimulus presentations occurred in a random order with an average inter-trial interval (ITI) of 3 min (range: 120 s-240 s). Rats remained in the chamber for 2 min following the final stimulus presentation.

##### Phase 2 Extinction

Following conditioning, rats were matched on their conditioned responding to the target stimulus across conditioning and were assigned to either an extinction or no-extinction control condition. On day 5, two minutes following placement in the conditioning chambers, rats in the extinction condition received twenty non-reinforced presentations of the target stimulus, whereas rats in the control condition received twenty presentations of the control stimulus. For both conditions, the cue and ITI durations were identical to those described for Phase 1. Ninety minutes following the first stimulus presentation, rats were perfused and brains were collected for immunofluorescent staining for Fos, β-gal, PKCδ, and SOM expression.

#### Experiment 2: Extinction-activated neurons in the BLA are not critical for extinction expression

##### Magazine Training, Phase 1 Conditioning, Phase 2 Extinction

Identical to Experiment 1 unless otherwise specified.

##### Phase 2 Extinction

Ninety minutes following the first stimulus presentation (target or control), rats were infused with either Daun02 or vehicle into the BLA. Rats receiving Daun02 infusions following presentations of the control cue served as the control for the specificity of the Daun02 manipulation; that is, they served to confirm that deleting cells that are activated by other stimuli had no effect on the neuronal ensembles activated by the target stimulus. This yielded four groups: extinction-vehicle, extinction-Daun02, control-vehicle, and control-Daun02.

##### Test for Extinction

On day 8, three days after extinction training, rats were tested for responding to the target stimulus. Rats were placed in the conditioning context, and after a 2-min adaptation period, the target stimulus was presented twenty times, non-reinforced, with identical ITI to that described previously.

#### Experiment 3a: Extinction-activated neurons in the CN are critical for extinction expression

##### Magazine Training, Phase 1 Conditioning, Phase 2 Extinction, Test for Extinction

All phases were identical to those described for Experiment 2 with the exception that Daun02 and vehicle solutions were infused into the CN and unless otherwise specified.

##### Phase 2 Extinction

For rats in the reinforced control experiment, Phase 2 training consisted of 20 reinforced trials of the target stimulus with identical ITI and Daun02/vehicle infusion into the CN to that described above.

##### Test for Extinction

On day 8, three days after extinction training, rats were tested for responding to the target stimulus. Rats were placed in the conditioning context, and after a 2-min adaptation period, the target stimulus was presented twenty times, non-reinforced, with identical ITI to that described previously. Following this Test, a subset of rats from the extinction-vehicle and extinction-Daun02 conditions were kept and later tested for subsequent extinction learning (the next day) and spontaneous recovery (9 days later). The remaining rats were perfused for confirmation of the Daun02 manipulation.

#### Experiment 3b: The original extinction-activated neurons in the CN regulate subsequent extinction learning and spontaneous recovery

##### Test for Re-Extinction

Twenty-four hours following Test for Extinction, rats were placed back in the conditioning context and tested again for responding to the target stimulus. After a 2-min adaptation period, the target stimulus was presented twenty times, non-reinforced, with identical ITI to that described previously.

##### Test for *Spontaneous Recovery*

Eight days after Test for Re-Extinction, rats were tested for responding to the target stimulus in the same manner as previously described.

#### Experiment 4: Deletion of extinction-activated neurons in the CN enhances reinstatement

##### Magazine Training, Phase 1 Conditioning, Phase 2 Extinction, Test for Extinction

Identical to Experiment 1 with the exception that all rats underwent extinction training to the target stimulus during Phase 2 Extinction.

##### Reinstatement

On day 8, three days after extinction training, rats in the reinstatement condition were placed back in the conditioning context, and received unsignalled exposure to the sucrose pellets in a manner identical to that described for magazine training (40-min session). This session consisted of delivering a sucrose pellet into the magazine at two-minute intervals (total of 20 pellets). No stimuli were presented during this phase. Rats in the control condition were handled for 30 s in their home-cage.

##### Test for Extinction

Twenty-four hours following reinstatement (day 9), rats were tested for responding to the target stimulus in a manner identical to that described for Experiment 2.

### Histology

Ninety minutes following the first cue presentation during Extinction training (Experiment 1) or Test (Experiments 2-4), rats received a lethal dose of sodium pentobarbital diluted 1:1 with 0.9% sodium chloride (120mg/kg) and perfused transcardially with 100 ml of 0.1M phosphate buffered saline (PBS; pH 7.4), followed by 200 ml of cold 4% paraformaldehyde in PBS, pH 7.4. Brains were then extracted and stored in buffered 4% paraformaldehyde for 1 hr, followed by 20% sucrose in PBS for 24 hr and subsequently dried and stored at -80°C until slicing. Brains were sectioned coronally at 40 µm through the BLA or CN using a Thermo Cryotome FE cryostat. One in every four sections were collected and stored in 0.1% phosphate buffered azide at 4°C.

For cannula placement, one in every four sections were collected on a slide and stained with cresyl violet. The location of the cannulation tips was determined under a microscope using the boundaries defined by the atlas of Paxinos and Watson (1997). Rats with incorrect placements or infection were excluded from statistical analysis. The numbers of rats excluded from each experiment based on incorrect placements or infection were seven rats in Experiment 2, nine rats in Experiment 3 (two rats in the reinforced control experiment), and thirteen rats in Experiment 4. Two rats from Experiment 2 were identified as significant outliers using the Grubb’s outlier test (ps<0.05), one on Conditioning and one on Test for Extinction, and excluded from all statistical analyses. The final *n*s for each group are presented in the figure legend for each experiment.

#### X-gal histochemistry for β-gal

X-gal staining was used to visualise β-gal as an indicator of neuronal activation and verify the effectiveness of the Daun02 manipulation (Koya *et al*., 2009; Koya et al., 2016). Unless specified, standard laboratory chemicals were purchased from Bioshop and Sigma-Aldrich. Free-floating sections were washed three times for 10 min each in 0.1M PBS (pH 7.4) and incubated in an X-gal reaction solution containing 2.4 mM X-gal (XGA002), 100 mM sodium phosphate, 100 mM sodium chloride, 5 mM EGTA, 2 mM magnesium chloride (MgCl_2_), 0.2% Triton X-100, 5 mM potassium ferricyanide (K_3_FeCN_6_), 5 mM potassium ferrocyanide (K_4_FeCN_6_) at 37°C for 4-5 hr with gentle shaking. Sections were then washed three times for 10 min each in 0.1M PBS (pH 7.4), mounted onto 4% gelatin-coated slides and air-dried for 24 hr. Slides were then dehydrated in increasing concentrations of ethanol (70%, 95%, 100%), cleared in a d-Limonene-based solvent (CitriSolv, Decon Labs), and coverslipped with Permount (Fisher Scientific). Images of the BLA and CN were assessed using a bright field microscope (Carl Zeiss Microscopy) under 5X magnification. The total number of β-gal positive cells were quantified manually by two observers in a blind manner by sampling 200 µm around the injection site from three to seven sections per region per rat. The mean correlation between the two scores were high (Pearson r = 0.89). Each rat represented a single observation for each brain area in the figures and statistical analyses.

#### Immunofluorescence double-labelling histochemistry for Fos/β-gal, Fos/PKCδ or Fos/SOM

We used double-label fluorescent immunohistochemistry to characterize BLA and CN neurons activated during extinction training. For Fos/β-gal double-labelling, free-floating sections were washed three times for 10 min each in 0.01M PBS (pH 7.4) and incubated in 10% normal goat serum (NGS, VectorLabs, S-1000) diluted in PBS containing 0.5% Triton X-100 for 2 hr at room temperature. The sections then incubated for 48 hr at 4°C in rabbit anti-c-Fos (1:1000, Cell Signalling Technology, 2250S, RRID: AB_2247211) and chicken anti-beta galactosidase (1:1000, Abcam, ab9361, RRID: AB_307210) diluted in PBS containing 0.5% Triton X-100 and 2% NGS. Sections were then washed three times in PBS (10 min each) and incubated in goat anti-rabbit Alexa Fluor 488 (1:200, Invitrogen, A-11034, RRID: AB_2576217) and goat anti-chicken Alexa Fluor 568 (1:200, Invitrogen, A-11041, RRID: AB_2534098) diluted in PBS containing 0.5% Triton X-100 and 2% NGS for 4 hr at room temperature. The sections were then rinsed three times in PBS (10 min each) and mounted onto 4% gelatin-coated slides. Slides were air-dried for 24 hr and coverslipped with Mowiol.

For Fos/PKCδ/SOM labelling, sections were washed three times for 10 min each in 0.01M PBS (pH 7.4) and incubated in 10% normal horse serum (NHS, Jackson Immuno Research Laboratories, 008-000-121) diluted in PBS containing 0.5% Triton X-100 for 2 hr at room temperature. The sections then incubated for 48 hr at 4°C in rabbit anti-c-Fos (1:1000, Cell Signalling Technology, 2250S, RRID: AB_2247211), mouse anti-PKCδ (1:1000, BD Biosciences, 610398, RRID: AB_2534098), and rat anti-somatostatin primary antibody (1:1000, Millipore Sigma, MAB354) diluted in PBS containing 0.5% Triton X-100 and 2% NHS. Sections were then washed three times in PBS (10 min each) and incubated in donkey anti-rabbit Alexa Fluor 594 (1:500, Jackson ImmunoResearch Laboratories, 711-585-152, RRID: AB_2340621), donkey anti-mouse Alexa Fluor 488 (1:500, Jackson ImmunoResearch Laboratories, 715-545-150, RRID: AB_2340846), and donkey anti-rat Alexa Fluor 647 (1:500, Jackson ImmunoResearch Laboratories, 712-605-153, RRID: AB_2340694) diluted in PBS containing 0.5% Triton X-100 and 2% NHS for 4 hr at room temperature. The sections were then rinsed three times in PBS (10 min each), mounted onto 4% gelatin-coated slides, air-dried for 24 hr, and coverslipped with Mowiol.

All fluorescent images of BLA and CN were captured using an epifluorescent microscope (Nikon TI) at 20X magnification and quantified using ImageJ software. The total number of cells positive for Fos and co-labelled with β-gal, PKCδ or SOM from the BLA and CN (specifically the lateral and capsular subdivision of the CN) were counted manually and using an automated counting script on ImageJ in a blind manner. The mean correlation between the two scores was high (Pearson r = 0.88). Each rat represented a single observation for each brain region in the statistical analyses and figures.

### Data analysis

Percent time spent in the magazine was used to measure conditioned responding following the onset of each cue presentation. Data were analyzed in SPSS 24.0 (IBM, New York, USA) using two-way repeated measures analysis of variance (ANOVA). Where appropriate, adjustments with Bonferroni were made for multiple comparisons. Neuronal counts were analyzed using independent samples t-tests. The percentage of overlap with Fos-positive neurons are reported as mean ± SEM. Significance was set at α = 0.05. For repeated measures ANOVA, Greenhouse–Geisser sphericity corrections were used when ε < 0.75. Standardized confidence intervals (CIs; 95% for the mean difference) and measures of effect size (η_p_^2^ for ANOVA and Cohen’s *d* for contrasts; see Cohen, 1988) are reported for each significant comparison. Data for each experiment and phase were reported and analyzed using all trials with the exception of Test for Reinstatement in Experiment 4, where reinstatement differences were present at the start of the test (first 6 trials) and subsequently masked by the effect of non-reinforcement. Therefore, in addition to using all trials, data for the first 6 trials were also reported.

## ACKNOWLEDGEMENTS

We would like to thank Dr. Shaun Khoo for assistance with cell counting, Dr Douglas Funk for initial input on Daun02 sourcing, and Dr. Rosemary Bagot for insightful discussions and invaluable comments on an earlier draft of our manuscript. This work was supported by FRQNT Nouveaux Chercheurs grant (MDI; 2017-NC-198182); a NARSAD Young Investigator grant (MDI; Grant # 24748); a CIHR Project Grant (MDI; Grant # PJT-155927); the Canada Research Chairs program (MDI; Grant # 950-230456); a Concordia University Horizon post-doctoral fellowship (BPPL); and a FRQS post-doctoral fellowship (to BPPL). The authors report no conflict of interest. All correspondence to be addressed to Dr. Mihaela Iordanova.

## Supplemental Figure Legend

*Supplemental Figure 1. CN but not BLA neurons are preferentially recruited during extinction*. Mean (+SEM) cell counts per mm2 for PKCδ or SOM positive neurons, Fos+PKCδ or Fos+SOM double-labelled neurons during extinction (top) and control (bottom) in the A) BLA and B) CN.

