## Supplemental for "A critical contribution of a sparse neuronal ensemble in the amygdala central nucleus to extinction"

Supplemental Information:

Results

**SUPPLEMENTAL RESULTS**

**Experiment 1: CN but not BLA neurons are preferentially recruited by extinction**

*Phase Conditioning 1.* Conditioned approach was acquired to the reinforced target, but not the non-reinforced control cue across Conditioning. A mixed ANOVA revealed no main effect of group (*F*_(1, 10)_ = 0.20, *p =* 0.66, 95% CI [-0.61, 0.40]), a main effect of cue (*F*_(1, 10)_ = 157.90, *p* < 0.001,
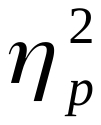
 = 0.94, 95% CI [2.87, 4.10]), and no group x cue interaction (*F*_(1, 10)_ = 0.17, *p* = 0.69,
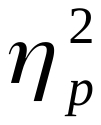
 = 0.017, 95% CI [-1.47, 1.01]), indicating that the increase in conditioned responding was specific only to the reinforced cue in all groups.

*Phase 2 Extinction.* Conditioning was followed by extinction training in half of the cohort. Extinction consisted of non-reinforced presentations of the target cue. In the remaining half of the cohort, the control cue was presented non-reinforced. A mixed ANOVA revealed a main effect of group (*F*_(1, 10)_ = 20.99, *p* = 0.001, 95% CI [0.55, 1.59], d = 2.64), trial (*F*_(5.34, 53.40)_ = 2.40, *p* = 0.046, ε = 0.28,
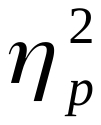
 = 0.19), and a trial x group interaction (*F*_(5.34, 53.40)_ = 2.36, *p* = 0.049, ε = 0.28,
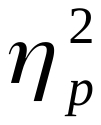
 = 0.19), indicating that a reduction in responding was evident in the extinction group.

Quantification of PKCδ-positive neurons within the BLA revealed no differences between the extinction and control groups (t(10) = 0.70, p = 0.50, 95% CI [-1.69, 0.88]). Similarly, there were no differences between the number of Fos + PKCδ double-labelled neurons (t(10) = 0.73, p = 0.49, 95% CI [-1.71, 0.87]), SOM-positive neurons (t(10) = 0.34, p = 0.74, 95% CI [-1.48, 1.09]), nor Fos + SOM double-labelled neurons (t(10) = 0.20, p = 0.85, 95% CI [-1.17, 1.40]; Supplemental Figure 1A).

Within the CN, rats in the control group exhibited more PKCδ-positive neurons compared to extinction rats (t(10) = 4.06, p = 0.002, 95% CI [-3.63, -1.06]). However, there were no differences between the groups in the number of Fos + PKCδ double-labelled neurons (t(10) = 0.092, p = 0.93, 95% CI [-1.34, 1.23]), SOM-positive neurons (t(10) = 0.32, p = 0.76, 95% CI [-1.47, 1.10]), and Fos + SOM double-labelled neurons (t(10) = 1.12, p = 0.31, 95% CI [-0.64, 1.93]; Supplemental Figure 1B).

**Experiment 2: Extinction-activated neurons in the BLA are not critical for extinction expression**

*Phase 1 Conditioning*. Conditioned approach was acquired to the target but not the control cue. A mixed ANOVA revealed no main effect of group (*F*_(1, 31)_ = 0.62, *p =* 0.44, 95% CI [-0.73, 0.39]), a main effect of cue (*F*_(1, 31)_ = 114.71, *p* < 0.001,
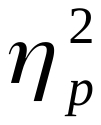
 = 0.79, 95% CI [1.42, 2.09]), and no group x cue interaction (*F*_(1, 31)_ = 0.11, *p* = 0.74,
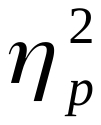
 = 0.004, 95% CI [-0.94, 0.72]), indicating that acquisition of conditioned responding was specific to the reinforced cue for all groups.

*Phase 2 Extinction*. Following conditioning, rats received non-reinforced presentations of either the target or the control cue during extinction. A mixed ANOVA revealed a main effect of group (*F*_(1, 31)_ = 26.29, *p* < 0.001,
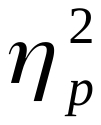
= 0.46, 95% CI [0.46, 1.54], d = 1.88), trial (*F*_(9.18, 284.49)_ = 8.90, *p* < 0.001, ε = 0.48,
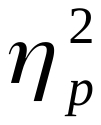
 = 0.22), and a trial x group interaction (*F*_(9.18, 284.49)_ = 4.54, *p* < 0.001, ε = 0.48,
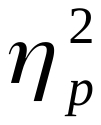
 = 0.13), indicating a reduction in responding across trials in the extinction groups. This training was followed by Daun02 or vehicle infusions into the BLA and all rats were tested three days later. Test statistics are reported in the main text.

**Experiment 3:** **Extinction-activated neurons in the CN are critical for extinction expression**

*Phase 1 Conditioning*. Conditioned approach was acquired to the target but not the control cue across Conditioning. A mixed ANOVA revealed no main effect of group (*F*_(1, 57)_ = 0.29, *p =* 0.59, 95% CI [-0.45, 0.29]), a main effect of cue (*F*_(1, 57)_ = 410.04, *p* < 0.001,
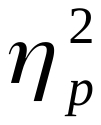
 = 0.88, 95% CI [2.04, 2.49]), and no group x cue interaction (*F*_(1,57)_ = 0.001, *p* = 0.98, ε = 0.74,
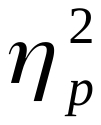
 < 0.001, 95% CI [-0.54, 0.56]), indicating that acquisition was specific only to the reinforced cue for all groups.

*Phase 2 Extinction*. Following conditioning, rats received non-reinforced presentations of either the target or the control cue during extinction. A mixed ANOVA revealed a main effect of group (*F*_(1, 57)_ = 47.39, *p* < 0.001,
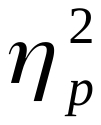
= 0.45, 95% CI [0.59, 1.25], d = 2.07), trial (*F*_(12.82, 730.59)_ = 5.57, *p* < 0.001, ε =0.68,
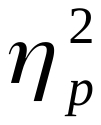
 = 0.089), and a trial x group interaction (*F*_(12.82, 730.59)_ = 3.48, *p* < 0.001, ε = 0.68,
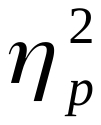
 = 0.057), indicating a reduction in responding across trials in the extinction groups. This training was followed by Daun02 or vehicle infusions into the CN.

*Test*. All rats were then tested for conditioned responding to the target cue three days later. A mixed ANOVA revealed a main effect of training (*F*_(1, 57)_ = 5.94, *p* = 0.018,
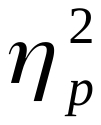
= 0.094, 95% CI [-0.67, 0], d = 0.62) and drug (*F*_(1, 57)_ = 6.63, *p* = 0.013,
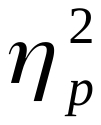
= 0.10, 95% CI [0.02, 0.69], d = 0.76), but no training x drug interaction (*F*_(1, 57)_ = 0.88, *p* = 0.35, 95% CI [-0.21, 0.47]). Post-hoc analyses are reported in the main text.

**Experiment 4: Deletion of extinction-activated neurons in the CN enhances reinstatement**

*Phase 1 Conditioning*. Conditioned approach was acquired to the target but not the control cue across Conditioning. A mixed ANOVA revealed no main effect of group (*F*_(2, 22)_ = 0.13, *p =* 0.88), a main effect of cue (*F*_(1, 22)_ = 147.30, *p* < 0.001,
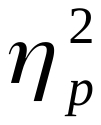
 = 0.87, 95% CI [-2.64, -1.71]), and no group x cue interaction (*F*_(2, 22)_ = 0.45, *p* = 0.65), indicating that acquisition was specific to the reinforced cue for all groups.

*Phase 2 Extinction*. Non-reinforced presentations of the target cue during extinction revealed a reduction in responding to the target cue. A mixed ANOVA revealed no main effect of group (*F*_(2, 22)_ = 2.36, *p* = 0.12), a main effect of trial (*F*_(3, 66)_ = 5.62, *p* = 0.002,
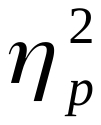
 = 0.20), and no trial x group interaction (*F*_(2, 22)_ = 0.48, *p* = 0.63) indicating that the rate of reduction in responding was similar for all groups. This training was followed by Daun02 or vehicle infusions into the CN.

*Test*. On Test, unsignalled exposure to pellets the day before reinstated responding to the extinguished target cue in the vehicle group compared to the controls during the beginning of the session (i.e. first-six trials, statistics reported in the main text) as expected, but these early differences were masked by non reinforcement across the entire test (across all trials, *F*_(1, 22)_ = 3.54, *p* = 0.22, 95% CI [-0.14, 0.90]). The rest of the comparisons were reported in the main text.
